# Plant Metabolic Network: A multi-species resource of plant metabolic information

**DOI:** 10.1101/2021.03.30.437738

**Authors:** Charles Hawkins, Daniel Ginzburg, Kangmei Zhao, William Dwyer, Bo Xue, Angela Xu, Selena Rice, Benjamin Cole, Suzanne Paley, Peter Karp, Seung Yon Rhee

## Abstract

Plant metabolism is a pillar of our ecosystem, food security, and economy. To understand and engineer plant metabolism, we first need a comprehensive and accurate annotation of all metabolic information across plant species. As a step towards this goal, we previously created the Plant Metabolic Network (PMN), an online resource of curated and computationally predicted information about the enzymes, compounds, reactions, and pathways that make up plant metabolism. Here we report PMN 15, which contains genome-scale metabolic pathway databases of 126 algal and plant genomes, ranging from model organisms to crops to medicinal plants, and new tools for analyzing and viewing metabolism information across species and integrating omics data in a metabolic context. We systematically evaluated the quality of the databases, which revealed that our semi-automated validation pipeline dramatically improves the quality. We then compared the metabolic content across the 126 organisms using multiple correspondence analysis and found that Brassicaceae, Poaceae, and Chlorophyta appeared as metabolically distinct groups. To demonstrate the utility of this resource, we used recently published sorghum transcriptomics data to discover previously unreported trends of metabolism underlying drought tolerance. We also used single-cell transcriptomics data from the *Arabidopsis* root to infer cell-type specific metabolic pathways. This work shows the continued growth and refinement of the PMN resource and demonstrates its wide-ranging utility in integrating metabolism with other areas of plant biology.

**One-sentence Summary:** The Plant Metabolic Network is a collection of databases containing experimentally-supported and predicted information about plant metabolism spanning many species.

## Introduction

Plant compounds are critical for the health, growth, and development of not only the plant, but also our planet and its biosphere. They allow the plant to defend itself from biotic and abiotic stressors (Weng 2014). The products of plant metabolism are also critical for humans, being the source of most human nutrition and numerous medicinally-useful compounds (Wurtzel and Kutchan 2016). It is therefore critical that we can understand, predict, and influence plant metabolism for the furtherance of economic, public health, and environmental preservation goals.

To provide the research community with comprehensive information about plant small-molecule metabolism, we previously introduced the Plant Metabolic Network (PMN), a plant-specific online resource of metabolic databases (Schläpfer et al. 2017). Accessible at https://plantcyc.org, the resource contains known plant metabolites, the reactions that create and consume them, the enzymes that catalyze the reactions, and the pathways into which the reactions can be organized. PMN consists of PlantCyc, a database of all experimentally-supported information found in the literature from any plant species, as well as single-species databases with a mix of experimentally-supported and computationally-predicted information, which allow researchers to explore each species’ unique metabolism.

The single-species databases were created using a computational pipeline we developed (Schläpfer et al. 2017). This pipeline is organized into three major stages: Enzyme prediction, done with the Ensemble Enzyme Prediction Pipeline (E2P2) software (Chae et al. 2014; Schläpfer et al. 2017); pathway, reaction, and compound prediction, done with the PathoLogic software (Karp et al. 2011; Karp et al. 2016; Karp et al. 2019); and pathway refinement, done with the Semi-Automated Validation Infrastructure (SAVI) software (Schläpfer et al. 2017). E2P2 predicts enzymatic functions of the proteins in a plant’s genome based on a reference protein sequence dataset (RPSD) using BLAST (Altschul et al. 1990) and PRIAM (Claudel-Renard et al. 2003). PathoLogic, distributed as part of the Pathway Tools software (Karp et al. 2019), takes in the enzyme annotation and retrieves from MetaCyc (Caspi et al., 2019), a pan-species reference database of metabolism that serves as a reference for PMN, all reactions that E2P2 predicted to be catalyzed by those enzymes, and predicts pathways based on the reaction complement (Schläpfer et al. 2017). Finally, SAVI applies previous pathway-level curation decisions to the new database. For example, a pathway might have been marked by curators to be present in all plants, in which case the pathway, along with its reactions and compounds, will be added to any plant database for which it was not predicted by PathoLogic, though the pathway will not have any enzymes associated to it. This pipeline enables the creation of a genome-scale metabolic pathway database for any plant species with a sequenced genome or transcriptome.

Here we describe PMN 15, the latest release of PMN that has grown substantially in both content and tools. We demonstrate the utility of the PMN resource by applying recently published omics data to gain insights into plant physiology and cellular level metabolism. Additionally, we systematically compare 126 species in the context of metabolism to identify metabolic domains and pathways that distinguish plant families. Finally, we present new website tools for viewing and analyzing metabolic data including a Co-Expression Viewer and subcellular boundaries for metabolic pathways.

## Results

### PMN is a comprehensive resource of plant metabolism databases

PMN is a collection of databases for plant metabolism with a substantial amount of experimentally supported information. The latest release (version 15) contains 126 databases of organism-specific genome-scale information of small-molecule metabolism alongside the pan-plant reference database PlantCyc (Figure 1). Together, these databases include 1,280 pathways, of which 1,163 have direct experimental evidence of presence in at least one plant species. In addition, PMN 15 includes 1,167,691 proteins encoding metabolic enzymes and transporters where 3,436 have direct experimental evidence for at least one assigned enzymatic function. There are 9,129 reactions (of which 34% have at least one enzyme from a plant species that has direct experimental evidence of catalyzing it), and 7,316 compounds. This large volume of metabolic information makes PMN a unique resource for plant metabolism.

**Figure 1.**
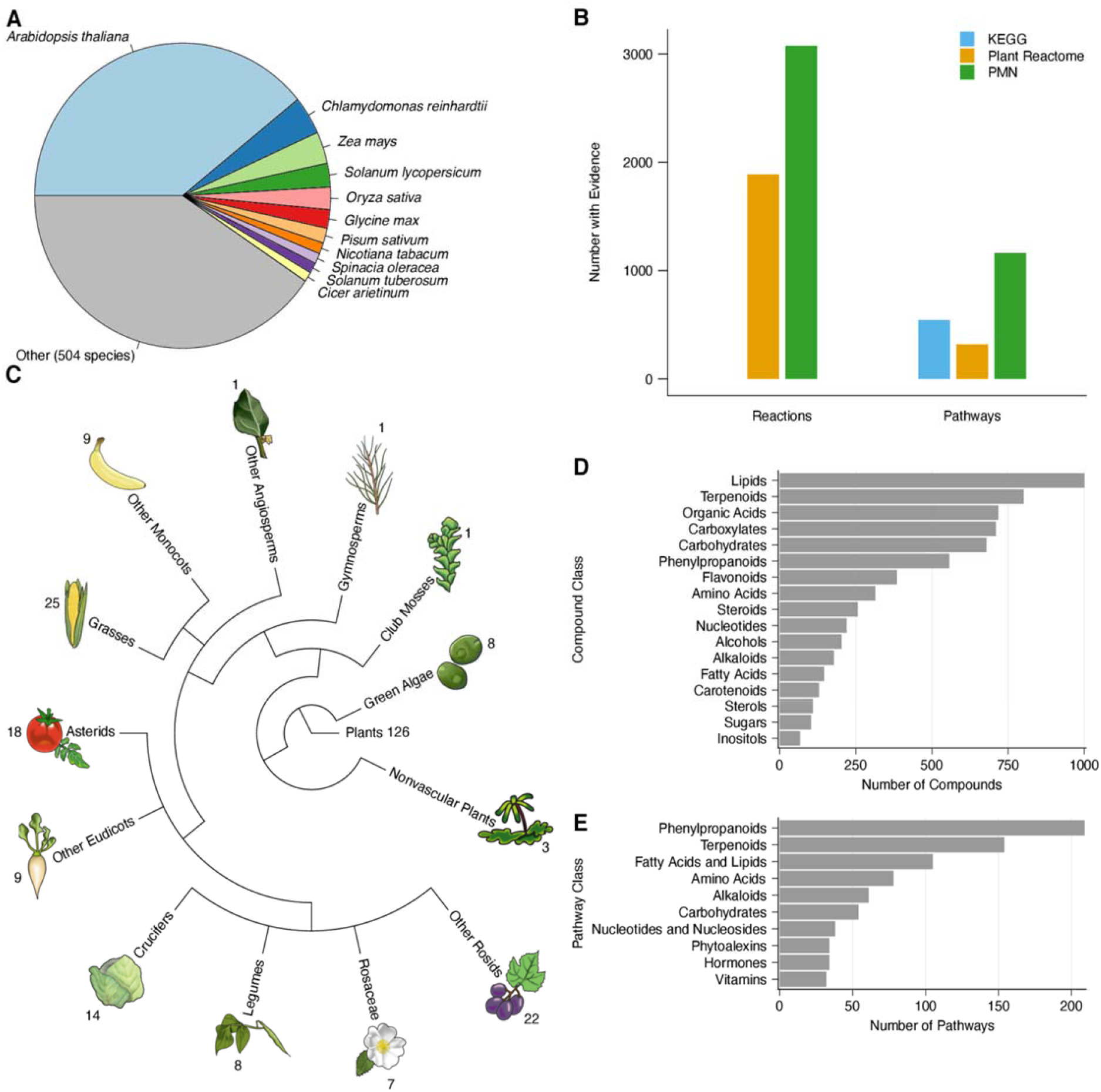
Plant Metabolic Network (PMN) content. The current content of PMN 15 and comparison to other databases. (A) Distribution of species based on the number of enzymes with experimentally supported evidence in PlantCyc. (B) Comparison of experimentally supported reaction and pathway data in PlantCyc, KEGG, and Plant Reactome databases. Evidence information for KEGG reactions was not accessible as of writing. (C) 126 species in PMN by phylogenetic group. (D – E) Distribution of the 7,316 compounds (D) and 1,280 pathways (E) in PMN 15 (PlantCyc + 126 species-specific databases), by class. The classes were manually selected from PMN’s class ontology. The classes are not exclusive; one compound or pathway may belong to multiple classes.

The reference database, PlantCyc, is a comprehensive plant metabolic pathway database. PlantCyc 15.0.1 contains experimentally supported metabolic information from 515 species. Most of the data come from a few model and crop species (Figure 1A). For example, *Arabidopsis thaliana* contributes to 43.4% of experimentally supported enzyme information in PlantCyc, followed by 7.46% from *Chlamydomonas reinhardtii* and 3.37% from *Zea mays*. Compared to other metabolic pathway databases such as KEGG (Kanehisa and Goto 2000; Kanehisa et al. 2017) and Plant Reactome (Naithani et al. 2017; Naithani et al. 2020), PlantCyc has substantially higher numbers of experimentally supported reaction and pathway data (Figure 1B). PlantCyc 15 includes 3,077 experimentally validated reactions with at least one curated enzyme and 1,163 curated pathways. Plant Reactome (Naithani et al. 2020) includes 1,887 and 320 curated reactions and pathways (Gramene release #61), while KEGG includes 543 experimentally-supported pathways as of February, 2021. The reference information in PlantCyc is incorporated into MetaCyc, which also includes experimentally supported metabolic information from non-plant organisms and is used to predict species-specific pathway databases (Caspi et al. 2020).

In addition to the reference database PlantCyc, PMN 15 contains 126 organism-specific metabolism databases (Figure 1C, Supplemental Table S1). These databases range widely in the plant lineage including several green algae and nonvascular plants. The majority of the plants are angiosperms with the Poaceae family most highly represented with 25 organisms. There are also 8 pairs of wild and domesticated plants, including rice, wheat, tomato, switchgrass, millet, rose, cabbage, and banana, alongside their wild relatives (Supplemental Table S2). Finally, PMN 15 includes 6 medicinal plants (species whose primary use is considered medicinal): *Camptotheca acuminata, Cannabis sativa, Catharanthus roseus, Ginkgo biloba, Salvia miltiorrhiza*, and *Senna tora*. The newest addition to the list of the medicinal plants is *Senna tora*, which is a rich source for anthraquinones and whose recent genome sequencing and metabolic complement annotation helped discover the first plant gene encoding a type III polyketide synthase catalyzing the first committed step in anthraquinone biosynthesis (Kang et al. 2020). This rich collection of species-specific metabolic pathway databases enables a wide range of analyses and comparisons.

PMN has grown significantly since its initial release (Figure S1A-H), with PMN 15 containing 2.5-fold more pathways, 4-fold more reactions, 3-fold more compounds, and 153-fold more enzymes than PMN 1. The focus on small-molecule metabolism means that processes involving the polymerization of macromolecules, such as transcription, translation, and DNA replication are excluded. Data in the PMN databases are represented using structured ontologies consisting of hierarchical classes to which pathways and compounds are assigned by PMN curators, which makes statistical enrichment analyses possible. The pathway and compound ontology classes, alongside the phylogeny of the included species, illustrate the breadth of metabolic information and species included in the database (Figure 1D, E). Prominent specialized metabolism classes such as terpenoids and phenylpropanoids are highly represented in the databases.

To promote interoperability and cross-referencing with other databases, PMN databases contain links to several compound databases such as ChEBI (Chemical Entities of Biological Interest) (Hastings et al. 2016), PubChem (Kim et al. 2021), and KNApSAcK (Nakamura et al. 2014). ChEBI release 197 has 58,829 entries and serves as a primary source of compound structural information during curation into PMN databases. Within PMN, 65% (4,746) of compounds link to ChEBI. PubChem is another chemical database, containing over 270 million chemical entries as of March 2021, and 95% (6,982) of PMN compounds link to it. Linking to these chemical databases provides a more in-depth source of information on the compounds and their physical and chemical properties. In summary, PMN is a broad resource for plant metabolism and continues to be under active development and expansion.

### Manual validation of pathway predictions reveals the continued necessity of manual curation

PMN databases include a large amount of computationally-predicted data. Predicting pathways for many species allows us to evaluate the quality of the predictions quantitatively. To estimate the extent of incorrectly-predicted pathways in the PMN databases, and to measure the overall accuracy of the computational predictions, both alone and in conjunction with manual curation, we evaluated the prediction of 120 randomly-selected pathways (approximately 10% of the 1280 pathways in PMN) on both the released organism-specific databases (also called Pathway Genome Databases (PGDBs) in Pathway Tools) and naïve prediction versions generated using only computational prediction (see Methods). Biocurators evaluated the pathway assignments to the 126 organisms currently in PMN, and classified them as “Expected” (predicted phylogenetic range is consistent with information in the literature), “Broader” (predicted taxonomic range includes expected range but is too broad), “Narrower” (predicted taxonomic range is within expected range but is too narrow), or as Non-PMN Pathways (NPP, not known to be present in plants or algae) (Figure 2, Supplemental Tables S3, S4). In the naïve prediction databases, only 15% of selected pathways were predicted within the phylogenetic ranges expected from the literature, and 58% were NPPs. In the released PGDBs, however, 78% of evaluated pathways were predicted as expected. In addition to correcting the prediction for 94% of all NPPs of the surveyed pathways, incorporating curated information also reduced the percent of pathways predicted beyond their expected phylogenetic ranges from 13% to 4%. Thus, the application of phylogenetic information and manual curation drastically improves the quality of pathway prediction throughout PMN databases over the use of computational prediction alone.

**Figure 2.**
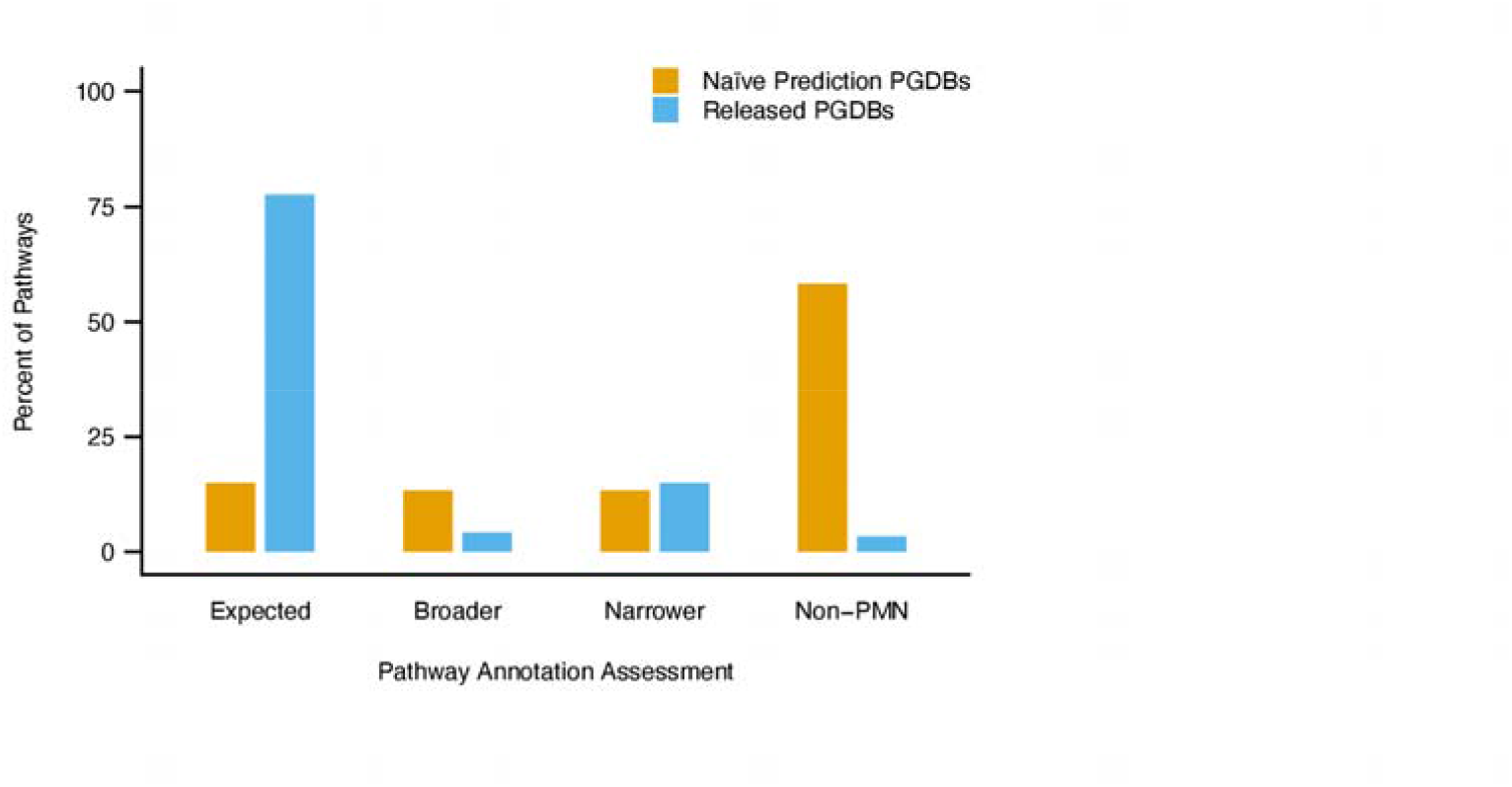
Manual pathway assessment. The result of manual review ofl20 randomly-selected pathways by biocurators. The plot shows the percentage of pathways in each assessment category in the naive prediction PGDB and the released PGDBs. Expected: Predicted and expected species are consistent; Broader: Pathway is predicted beyond its expected range; Narrower: Predicted range of the pathway is smaller than the expected range; Non-PMN: The pathway is not expected to be found in plants or green algae.

### PMN data can distinguish phylogenetic groups

PMN 15’s utility depends on the completeness and accuracy of the data it contains for its 126 organisms. Objectively evaluating the quality and richness of PMN’s data is not straightforward, however, because there is no “gold standard” to compare PMN against. If PMN 15 contains data that accurately reflect the diversity of all 126 organisms, it should be possible to differentiate known groups of plants based upon their metabolic data. If plants in a specific group cluster together based on their metabolic content, this may indicate that the unique metabolism of the group is well-represented in PMN. If a known group cannot be differentiated from others, this may indicate that more research and curation are needed to understand the group’s unique metabolism and can thereby guide future research and curation.

To determine whether different groups of plants can be differentiated solely by their metabolic capacity, we performed multiple correspondence analysis (MCA), a type of dimension reduction analysis that is similar to principal component analysis but can be used for categorical data (Tenenhaus and Young 1985; Greenacre et al. 2006). MCA was carried out using presence-absence matrices for pathways, reactions, and compounds (Figure 3 and Supplemental Figure S2; Supplemental Table S5). Reactions were considered present only if at least one enzyme in the species was annotated as catalyzing the reaction. Independently, the plants were categorized according to phylogenetic groups. Dimensions 1 and 3 of the pathway and compound MCA, and dimensions 1 and 2 of the reaction MCA, separated the species into several phylogenetic groups (Figure 3A and Supplemental Figure S2C, G, H). Phylogenetic groups that clearly cluster together and away from other groups include algae, non-flowering plants, Brassicaceae, and Poaceae (Figure 3A and Supplemental Figure S2G, H). Dimension 1 separates the chlorophytes from land plants and dimension 3 separates certain angiosperm families such as the Brassicaceae and Poaceae well. No clear separation was observed among other eudicot groups. In addition, dimension 2 of the pathway and compound MCA mostly separated a small number of highly curated species from all the rest (Figure S2A, E; Supplemental Table S5). Overall, the MCA clustering shows that some groups of plants can be readily differentiated based on their metabolic information (compounds, enzymes, reactions, pathways) in PMN, while other groups cannot, suggesting that further curation of species in these groups may be beneficial.

**Figure 3:**
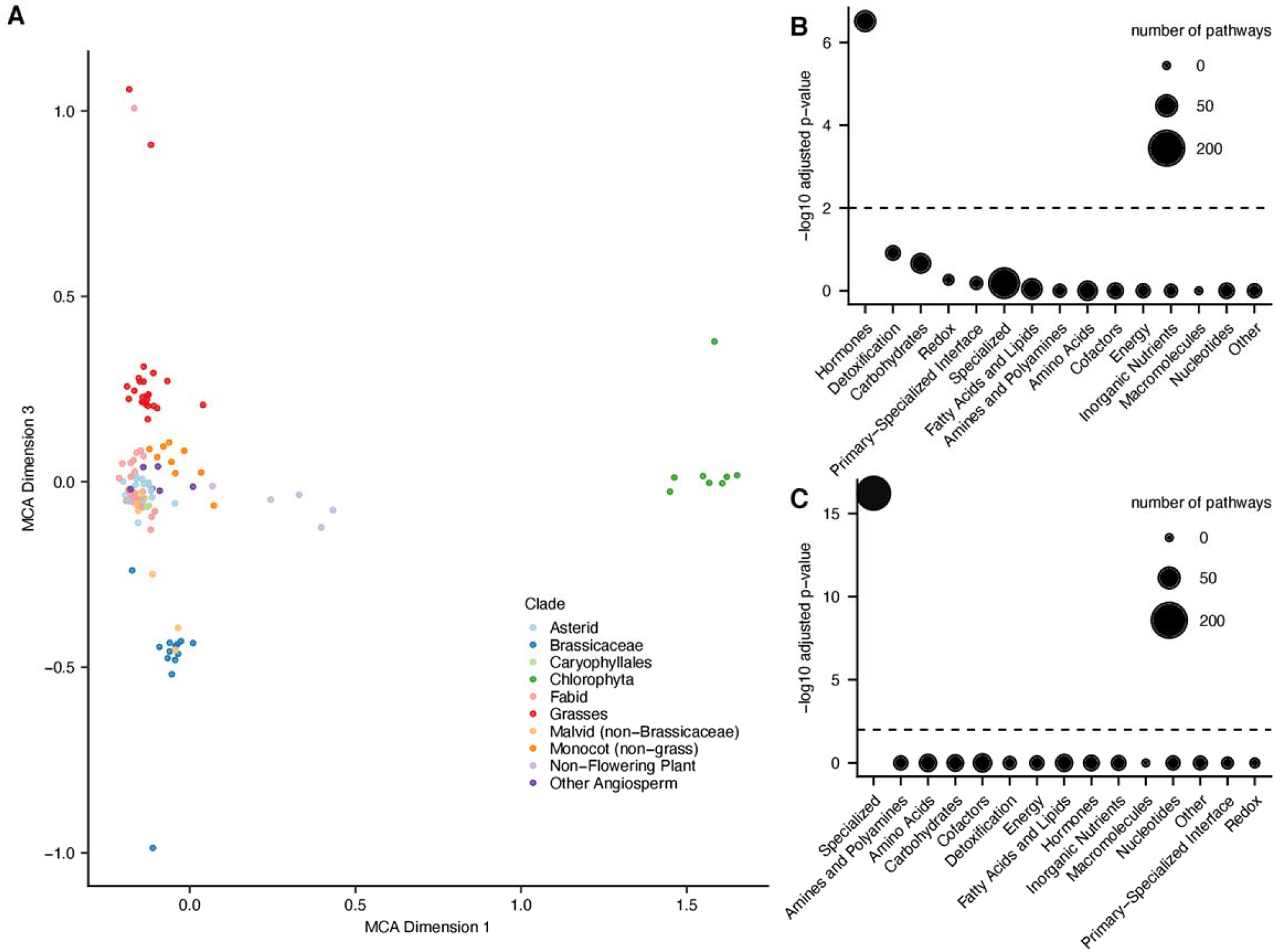
Pathway multiple correspondence analysis. Multiple correspondence analysis (MCA) performed on a binary matrix of pathway presence/absence in each species. (A) A scatter plot of dimensions 1 and 3 of the MCA; each dot is a plant species, and the coordinates are a dimensional reduction of the binary presence/absence vector of pathways. Color represents plant group information overlaid onto the plot. Brassicaceae, Poaceae, green algae, and nonseed plants are discernable as clusters. Dimensions 1 and 3 were selected because they illustrate the clustering well. (B) Metabolic domain enrichment for pathways explaining 95% of the variance in MCA dimensions 1 and 3; bubble size indicates the number of pathways meeting the 95% variance cutoff in each domain. Dashed lines represent p=0.01 significance threshold. P-values were corrected for multiple hypothesis testing at a false discovery rate (FDR) of 5%.

We next asked which metabolic pathways drive the separation of the taxonomic groups on each dimension (Supplemental Table S5). 70% of the variance in dimension 1 was described by 109 pathways, all of which were predicted to be either embryophyte-specific pathways or present in a larger proportion of embryophytes than chlorophytes. This mirrors the separation of the Chlorophyta cluster in dimension 1 of the MCA plot (Figure 3A; Supplemental Table S5). Similarly, 70% of the variance along dimension 3 was captured by 150 pathways, of which 81 were associated more strongly with Poaceae and 69 were associated more strongly with Brassicaceae (Figure 3A; Supplemental Table S5). The pathways that contributed 95% of the variance in dimension 1, which separates chlorophytes from embryophytes, were enriched for hormone metabolism (Figure 3B, adjusted p-value = 1.6E-07, hypergeometric test). Hormone metabolism may have helped support the increased complexity of land plants compared to their algal ancestors (Wang et al. 2015). In contrast, pathways responsible for clustering along dimension 3 were enriched for specialized metabolism (Figure 3C, adjusted p-value = 1.1E-22, hypergeometric test), which is more lineage-specific than other domains of metabolism and can help distinguish between clades of angiosperms (Chae et al. 2014). Thus, it appears that metabolic data in PMN can effectively differentiate groups of species not only by the presence or absence of specific pathways and reactions, but also by the types of metabolic processes which are related to their evolutionary divergence.

### Data analysis tools and applications with external datasets

PMN contains not only information about the compounds, reactions, and pathways of plant metabolism, but also a suite of tools to compare and analyze these data. For example, lists of pathways, reactions, compounds, genes, or other data objects can be assembled into SmartTables for further analyses, or to export data in a tabular format. Omics data, or any numeric data associated with genes, proteins, or compounds, can be overlaid onto the pathways and reactions associated with those genes, or uploaded into Pathway Tools’ Omics Dashboard (Paley et al. 2017; Paley et al., 2021), which allows users to visualize omics data across experimental timepoints and conditions at various scales of metabolism including broad metabolic domains, individual pathways, and genes. Here we demonstrate two applications of integrating omics data with PMN resources to gain novel insights about plant metabolism.

To demonstrate the utility of the Omics Dashboard in analyzing omics data within a metabolic context, we turned to a recently published transcriptomic survey of two sorghum cultivars, RTx430 and BTx642, subjected to drought stress at multiple points throughout the growing season (Varoquaux et al. 2019). RTx430 is tolerant to pre-flowering drought, whereas BTx642 is tolerant to post-flowering drought. To see if there was any difference in metabolic gene expression between the two cultivars in response to post-flowering drought, we examined differentially expressed genes (DEGs) in droughted plants compared to well-watered plants from the last week of watering (week 9 after sowing) to the first two weeks of post-flowering drought (weeks 10 – 11). We observed quantitative differences in global metabolic gene expression between the two cultivars, specifically the consistent down-regulation of biosynthetic activity from root tissues in the post-flowering drought sensitive cultivar RTx430 compared to relatively stable expression in the post-flowering drought tolerant cultivar BTx642 (Figure 4A). This observation is consistent with the authors’ findings that BTx642 demonstrated higher levels of redox balancing and likely experienced lower levels of reactive oxygen species stress, compared to RTx430, as a result of drought. By analyzing expression patterns of all metabolic genes, we observed widespread metabolic down-regulation in RTx430 root tissue, which was not reported previously (Varoquaux et al. 2019). To determine whether the consistent reduction of metabolic gene expression observed in RTx430 roots in response to drought was a global trend in the transcriptome or specific to metabolic genes, we compared relative expression levels of all non-metabolic root DEGs to all metabolic root DEGs in both cultivars during the same 3-week period. While the average relative expression decreased each week among both metabolic and non-metabolic genes in RTx430, the down-regulation was greater among metabolic genes at both time points (Supplemental Figure S3B). In contrast, BTx642 roots showed no difference in expression among both metabolic and non-metabolic genes in response to drought (Supplemental Figure S3B), suggesting a global metabolic homeostasis in sorghum drought tolerance. By comparing the *patterns* of expression among DEGs in root and leaf tissues, rather than solely the *number* of DEGs, analysis via the Omics Dashboards revealed that roots exhibited stronger genotype-specific responses to drought than leaves, which was not observed previously (Varoquaux et al. 2019). Drought-responsive DEGs were enriched in metabolic genes among both leaf (p = 2.2E-84, hypergeometric test) and root (p = 1.7E-114, hypergeometric test) tissues. However, contrary to the clear cultivar-specific trends shown in the root DEGs (Figure 4A), the metabolic genes did not show any clear trend in their expression patterns in the leaves of either cultivar as a result of drought (Figure S3A).

**Figure 4:**
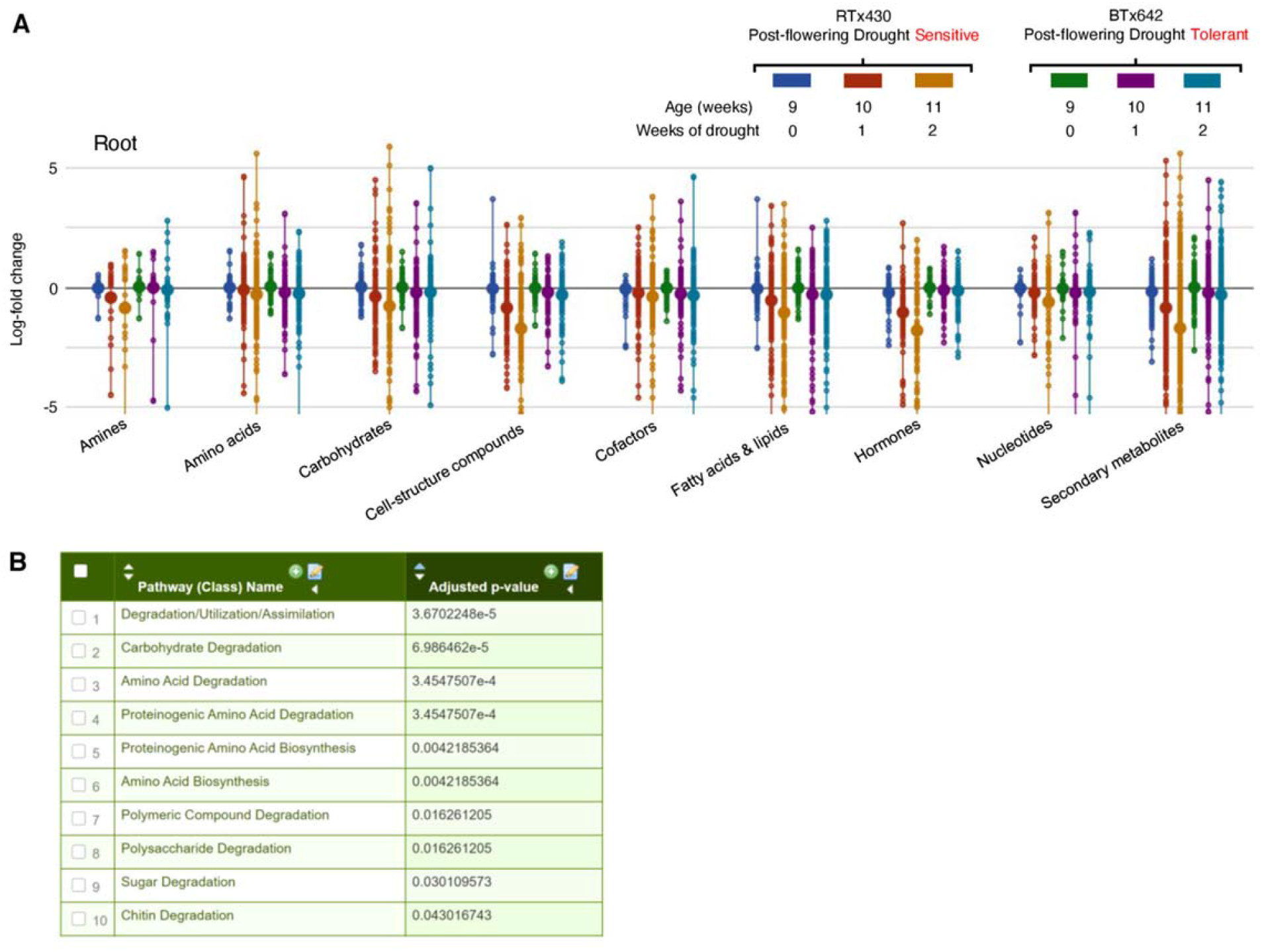
PMN’s Omics Dashboard and pathway enrichment analyses. (A) Omics Dashboard representation of global metabolic biosynthetic expression patterns among DEGs in response to drought in root tissues of two sorghum cultivars. The y-axis represents log_2_ fold-change between drought and non-drought conditions at each time point; positive values indicate higher expression under drought. Categories along the x-axis are major top-level pathway classes in Pathway Tools. (B) Screenshot of the results of a pathway enrichment performed in PMN, showing significantly enriched pathways among drought-responsive metabolic genes compared to all metabolic genes in RTx430 root tissue.

In addition to offering a visual overview of metabolism via the Omics Dashboard, PMN’s analytical toolkit allows researchers to easily conduct enrichment analyses among a set of genes or compounds of interest. From within a SmartTable, users can view the pathways associated with a set of genes or compounds, and can then ask whether those genes or compounds are enriched for specific pathways or classes of pathways. Broader metabolic classifications can also be added to the list of enriched pathways to better understand which area(s) of metabolism are most enriched. For example, among the set of drought-responsive DEGs in RTx430 roots, we observed an enrichment in various domains of carbohydrate and amino acid biosynthesis and degradation, in addition to chitin degradation, consistent with the authors’ observation of drought-induced responsiveness of biotic defense genes (Figure 4B). Thus, by combining PMN’s analytical capabilities with its broad set of metabolic data, users can find additional means of supporting existing hypotheses, uncovering novel insights, and finding new avenues for exploration in their own research.

The data-rich resources within PMN can also be integrated with other cutting-edge datasets to investigate novel biological questions. For example, single cell sequencing technologies, such as drop-seq and the 10X scRNA-Seq platform, have been adapted to plant cells to generate high-resolution transcriptomic profiles in *Arabidopsis* root cells (Denyer et al., 2019; Jean-Baptiste et al. 2019; Ryu et al., 2019; Shulse et al. 2019; Zhang et al., 2019; Wendrich et al., 2020). In this study, we downloaded and integrated datasets from five existing *Arabidopsis* root single-cell RNAseq studies to generate a comprehensive transcriptome profile (Supplemental Table S6). These single-cell level data allow us to investigate cell type specificity of metabolic pathways and domains at the transcript level. We define cell type-specific metabolic domains (or pathways) as those whose constituent genes show significantly higher expression levels (fold change ≥ 1.5, Wilcoxon test p-value 0.05) in certain cell types compared to their average expression level in total cells. Different metabolic domains showed overlapping as well as distinct cell type specificity (Figure 5A). First, epidermal and cortex cells were most metabolically active throughout the various domains of metabolism (Figure 5A). This is consistent with previous observations that the major groups of metabolites detected in *Arabidopsis* roots, including glucosinolates, phenylpropanoids, and dipeptides, were highly abundance in the cortex (Moussaieff et al. 2013). In contrast, maturing xylem showed relatively low metabolic activity as the major roles of these cells are structural support and water/soluble transport (Schuetz et al. 2013). Viewed from the level of metabolic domains, this analysis demonstrates a diverse range of metabolic activity across unique cell types in *Arabidopsis* roots.

**Figure 5:**
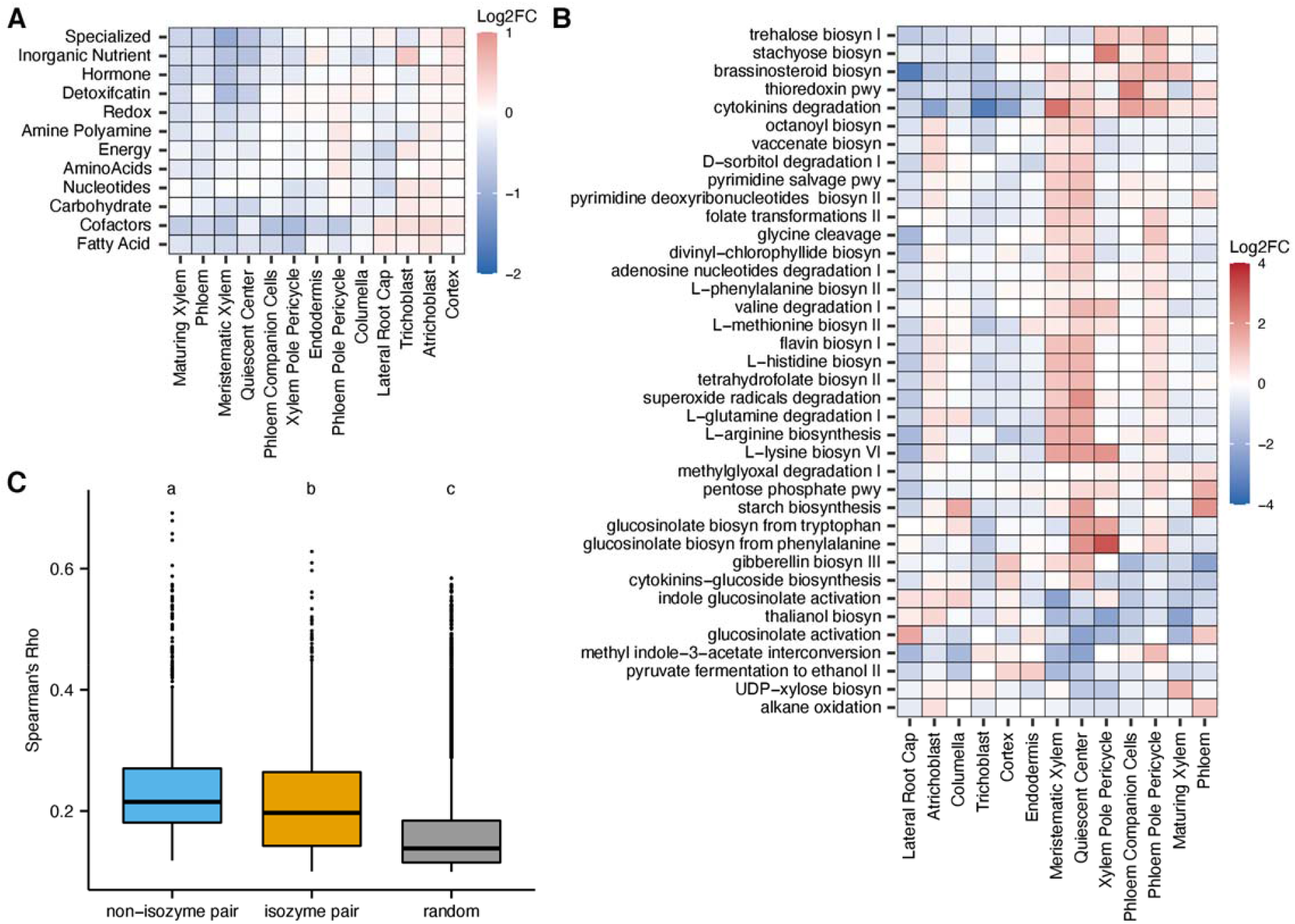
Comparison of metabolism across *Arabidopsis* root cell types. Cell type specificity of metabolic (A) domains and (B) pathways. Log2FC represents the log2 fold change of the expression level of a metabolic domain or pathway in a cell type over their average expression level in total cells. (C) Isozymes are more likely to be expressed in different cell types compared to other enzymes catalyzing different reactions in the same pathway. The box plot represents Spearman’s correlation coefficient computed to measure the gene expression pattern similarity between a pair of enzymes across *Arabidopsis* root cells. Letters above the boxes represent significantly different groups of p value < 0.05 as determined by one-way ANOVA followed by post-hoc Tukey’s test.

We next probed cell-type specificity of individual pathways. Among the 198 pathways associated with at least 10 genes, 40 pathways (20%) showed specificity in at least one cell type compared to their background gene expression levels represented by the average expression level of the pathway across all cell types (Figure 5B). For example, in actively dividing cells, such as meristematic xylem cells, pathways involved in pyrimidine, histidine, arginine, and lysine biosynthesis showed high activity (Figure 5B). These pathways are involved in essential metabolism, which are critical for maintaining cell division and growth. On the other hand, hormone biosynthesis pathways, such as cytokinin glucoside and gibberellin, showed high activity in the cortex. This is consistent with current understanding that the cortex is one of the predominant cell types that synthesizes these two hormones in the *Arabidopsis* root (Antoniadi et al. 2015; Barker et al. 2020). By elucidating cell type-level activity of metabolic pathways, we can begin to map metabolism at cellular and tissue levels, which will be instrumental in understanding how metabolism affects plant development and responses to the environment as well as enabling effective engineering strategies.

Similar to cell-type specificity, the concept of pathway divergence at the individual cell level can also be explored using single cell transcriptomics data. To probe this question, we asked whether isozymes catalyzing the same reaction are more likely to be expressed in different cells compared to enzymes catalyzing different reactions in the same pathway. Isozymes are defined as enzymes encoded by different genes catalyzing the same reaction, which are usually the result of gene duplication events. We computed Spearman’s correlation coefficient to measure gene expression pattern similarity between a pair of enzymes across *Arabidopsis* root cells. The coefficients computed based on single cell data were generally lower than that generated by bulk RNA-seq, which may be due to the sparseness of single cell transcriptomic profiles or high heterogeneity of gene expression across cells. Nonetheless, metabolic genes in the same pathway showed higher correlation than randomly sampled metabolic genes (Figure 5C), which suggests functional coordination between genes involved in the same pathway at the cellular level. Isozymes were much less correlated than enzyme pairs catalyzing different reactions in the same pathway. This indicates that isozymes may have evolved divergent expression patterns in root cells (Figure 5C). Since isozymes are often the results of gene duplication events, this diversified expression between isozymes may contribute to retaining duplicated genes through subfunctionalization or neofunctionalization and fine-tuning metabolic pathways at the cellular level (Panchy et al. 2016).

### New capabilities and integration with other databases

Recently we introduced the Pathway Co-Expression Viewer, which integrates information from PMN and ATTED-II (Obayashi et al. 2018), a database of gene co-expression, to visualize co-expression of the genes in a pathway for species represented in ATTED-II (*Arabidopsis thaliana, Glycine max* (soybean), *Solanum lycopersicum* (tomato), *Oryza sativa* (rice), *Zea mays* (maize), *Brassica rapa, Vitis vinifera* (grape), *Populus trichocarpa* (poplar), and *Medicago truncatula*). An example is shown in Figure 6A-B; Lysine biosynthesis is currently known to occur via two distinct routes, utilizing either diaminopimelate or α-aminoadipate as an intermediate. Its biosynthetic pathway in plants, cyanobacteria, and certain archaebacteria (PWY-5097) (Figure 6A) converts tetrahydrodipicolinate to L,L-diaminopimelate via L,L-diaminopimelate aminotransferase and is distinct from that of other prokaryotes and of fungi (Hudson et al. 2006). Lysine biosynthesis is of particular importance as it is both an essential amino acid not biosynthesized by mammals and it is the least abundant essential amino acid in cereals and legumes (Wang and Galili, 2016). The Pathway Co-Expression Viewer shows that the genes in this pathway exhibit high levels of co-expression. The co-expression levels of six pairs of genes are in the top 1% of co-expressed gene pairs within ATTED-II, while an additional 10 gene pairs are in the top 5% (Figure 6B, dark gray). This tool provides a convenient way of visualizing the co-expression of genes in a pathway and thus provides clues as to how the pathway may be regulated.

**Figure 6:**
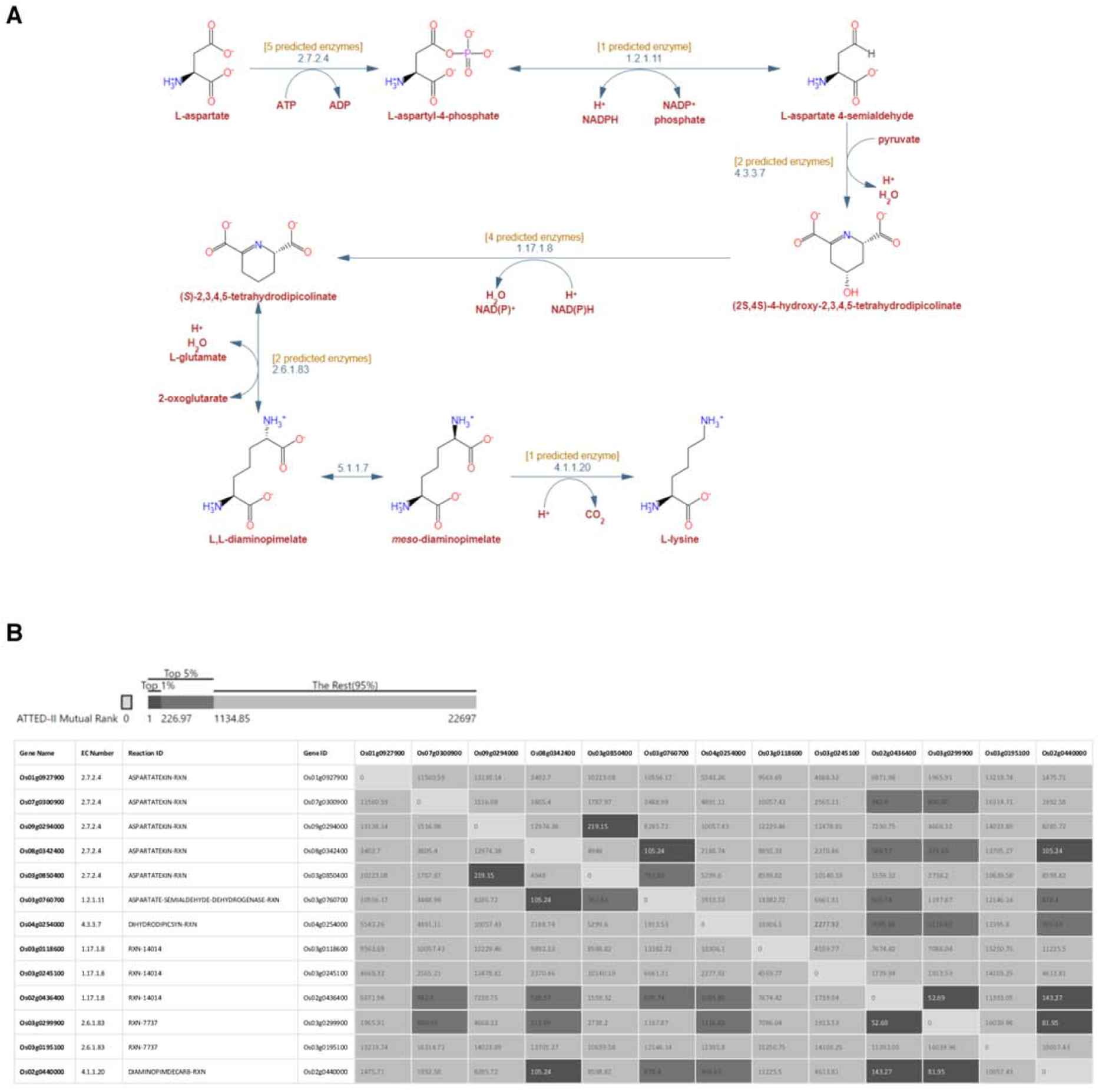
Pathway visualization tools. Exploration of the pathway visualization tools in PMN. (A) L-lysine biosynthesis VI (PWY-5097) in OryzaCyc 7.2.0. (B) co-expression view of the rice genes that code for enzymes which catalyze reactions in the lysine pathway. Genes are in both rows and columns. Each numerical cell shows the co-expression of the two genes as ATTED-II mutual rank (Obayashi et al. 2018). Lower numbers indicate stronger coexpression. Medium gray indicates the pairing is in the top 5% of the mutual rank score; dark gray indicates top 1%. Genes with no co-expression data have been manually removed from the table.

PMN 15 introduces an additional feature which provides a new way of visualizing pathways that span intracellular compartments and include transport reactions. For example, the glutamate-glutamine shuttle (PWY-7061; Figure 7) from AraCyc is a pathway in which glutamate and glutamine are exchanged between the mitochondria and chloroplast as a means of ridding the mitochondria of ammonium produced during photorespiration (Linka and Weber 2005). Membranes that separate compartments are rendered as gray bars, with both sides labelled, and transporters are shown as breaks in the gray bar with pairs of brown ovals on either side to suggest a channel. This new feature makes intracellular transport within pathways clearer and easier to visualize.

**Figure 7:**
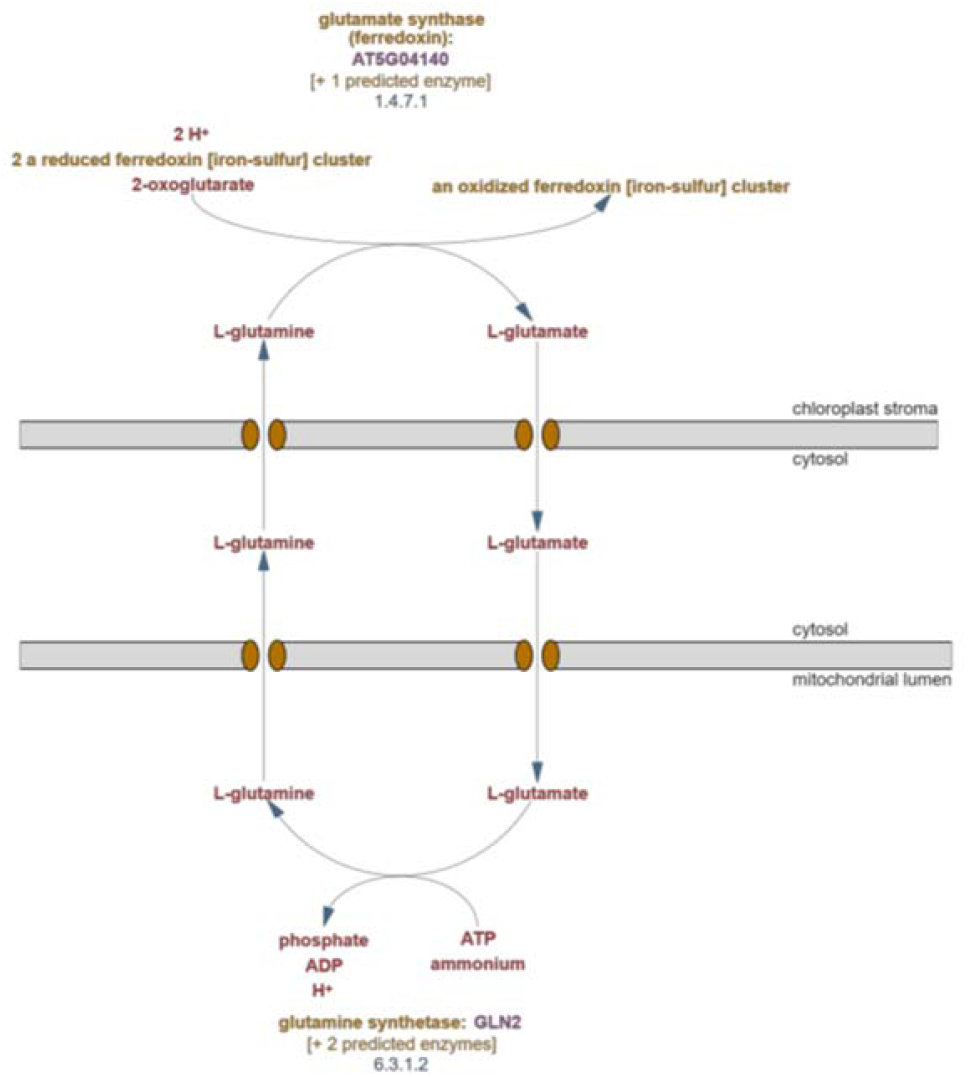
Visualization of pathway subcellular localization. Subcellular localization view introduced in PMN 15. The glutamate-glutamine shuttle (PWY-7061) in AraCyc 17.1.1 is shown, with chloroplast and mitochondrial outer membranes displayed on the diagram.

## Discussion

PMN 15 is an extensive and regularly-updated database of compounds, pathways, reactions, and enzymes for 126 plant and green algae species and subspecies as well as a pan-species reference database called PlantCyc. We examined the quality of the data contained in the databases by assessing the accuracy of pathway prediction via manual validation of a randomly-selected subset of predicted pathways. Using two publicly available transcriptomics datasets, we demonstrated how PMN resources can be leveraged to characterize and gain insights from omics data. The present work demonstrates that the Plant Metabolic Network can be a useful tool for various analyses of plant metabolism across species.

### Accuracy of PMN

The ability of PMN to enable research is dependent on the accuracy of its data. We therefore evaluated the performance of PMN’s metabolic reconstruction pipeline both in its entirety and using only computational prediction. The manual pathway validation revealed a number of pathways predicted to be present outside of their known taxonomic range, such as momilactone’s predicted presence across Poaceae despite being known to exist only in rice and a few other species, some outside of Poaceae (in which they appear to have evolved convergently) (Mao et al. 2020). While some of these results may reflect compounds that are, in fact, more widely distributed than currently thought, many such cases likely result from inaccurate prediction of enzymatic function by E2P2. The performance of enzyme function prediction using a sequence similarity approach can suffer when dealing with highly similar enzymes of a shared family (Schläpfer et al. 2017). In cases like momilactone, where the pipeline has predicted the pathway in species closely related to species known to possess it, it may be the case that the predicted species do have most of the enzymes necessary to catalyze the pathway, but that one or a few of the predicted enzymes actually have a different function *in vivo*. This may draw attention to cases where enzymes have gained new functions and allow for exploration of how enzymes evolve. Meanwhile, cases of universal plant pathways being predicted only in Brassicaceae may indicate the pitfalls of an overemphasis on *Arabidopsis* in curation and research, as key enzymes might be predicted less reliably outside of this clade. This might be the case if there are Brassicaceae-specific variations that may result in a failure to reliably predict orthologs. A focus on curating information from diverse species may improve the accuracy of the computational prediction, requiring less semi-automated curation to fix such errors.

Pathway misannotation in the naïve prediction pipeline (see Methods) could also be the result of PathoLogic’s incorrect integration of enzyme annotation with reference reactions. In addition to incorporating enzyme predictions, PathoLogic can infer pathways for a given species based on a number of additional considerations. For example, if a species contains an enzyme which catalyzes a reaction unique only to one pathway in the PGDB, the pathway is likely to be predicted to be present. Additionally, if all reactions of a pathway are predicted to be present, the pathway is likely to be predicted as. Using PathoLogic without taxonomic pruning thus provides increased prediction sensitivity while also increasing false positives (Karp et al. 2011; Schläpfer et al. 2017). By design, SAVI removes false-positive and adds false-negative pathways predicted by PathoLogic. Our analyses indicate that the predominant function of SAVI and PathoLogic’s taxonomic pruning currently is to remove false-positives and consequently restrict the taxonomic range of predicted pathways, consistent with previous analyses of SAVI’s performance (Figures 2, S2) (Schläpfer et al. 2017). Interestingly, our manual pathway assessment revealed that, in certain cases, SAVI should have increased the range of a predicted pathway and added it to more species than it was predicted for by PathoLogic. For example, the phytol salvage pathway (PWY-5107) is predicted to be present in all photosynthetic organisms (Valentin et al., 2006). While PathoLogic incorrectly restricted the predicted range of this pathway to include only angiosperms even without taxonomic pruning, SAVI did not correct this incorrect taxonomic restriction, nor did it assign the pathway to the few angiosperm species not predicted by PathoLogic to contain the pathway. Examples like this may represent errors in the manual curation decisions used by SAVI to make its correction, or it may reflect new information added to the literature after those curation decisions were made. Both possibilities represent important information in accurately representing metabolism across species and highlight the need to regularly update the curation rules upon which SAVI operates. We therefore reclassified the final pathway assignments in PMN 15 for each pathway whose classification after SAVI implementation was determined to be anything other than “Expected”. Through the continual process of introducing new species — and thus new pathways — into PMN, along with regular curation of those new pathway predictions, SAVI’s correction performance, and thus the overall value of data in PMN, should continue to improve over time.

### Other metabolic pathway databases

PMN strives to differentiate itself from other metabolic pathway databases through the quantity of curated and computational information, its comprehensive set of tools, and its specific focus on plants. Other, comparable databases include KEGG (the Kyoto Encyclopedia of Genes and Genomes) (Kanehisa and Goto 2000; Kanehisa et al. 2017; Kanehisa et al. 2019), Plant Reactome (Gramene Pathways) (Naithani et al. 2020), and WikiPathways (Slenter et al. 2018). Like PMN, these databases contain metabolic pathways along with their associated reactions, compounds, and enzymes. KEGG pathways represent broad metabolic reactions shared among many organisms, and it is common to map genes or compounds to KEGG pathways alongside Gene Ontology (GO) annotations for enrichment analyses. However, because KEGG pathways represent a generalized set of reactions leading to many possible compound classes (but not to specific compounds), it lacks the granularity to analyze metabolism on a species-specific level (Altman et al. 2013). For example, a recent study identified enriched KEGG pathways (e.g., “phenylpropanoid biosynthesis”) among genes belonging to gene families that were expanded in *Senna tora* compared with its relatives (Kang et al. 2020). Enrichment analysis of the same genes using PMN’s StoraCyc 1.0.0 identified individual phenylpropanoid biosynthetic pathways enriched among the gene set, such as coumarin biosynthesis. PMN and MetaCyc feature structured data that is both human-and machine-readable, making it possible for users to obtain pathway structure and other data for their own offline analysis and enabling features such as the pathway Co-Expression Viewer to be easily incorporated. WikiPathways is another pathway-centric database. WikiPathways is not plant-focused, and takes a crowd-sourced approach, in contrast with PMN’s focus on expert curation. Plant Reactome, another metabolism database, is specific to plants and green algae as PMN is. However, Plant Reactome uses *Oryza sativa* as a reference species to predict reactions and pathways to the 106 other species currently in the database and uses gene orthology to predict the presence of a pathway, where a pathway is predicted in a species if at least one rice ortholog for an enzyme in that pathway is present in that species (Naithani et al. 2020). Pathway prediction in PMN, on the other hand, is more stringent via its implementation through the PathoLogic and SAVI pipelines.

### Associations between metabolism and phylogeny

PMN is organized primarily by species, and a significant component of the expansion over its history has been in the form of adding new species and subspecies to it. In order for this to be a worthwhile endeavor and useful to the plant biology research community, the species databases need to be meaningfully differentiated from one another in ways that accurately reflect their metabolic differences. Multiple correspondence analysis was therefore performed to determine whether related species would cluster together, an indication that underlying biology is driving the differences in their database contents. The analysis revealed that some plant groups such as Brassicaceae, Poaceae, the green algae, and non-flowering plants each clustered together, showing that these major groups of plants can be readily differentiated based on their metabolic complements. Within the eudicots, however, there was little separation apart from the grouping of Brassicaceae. Other groups such as Rosaceae and Solanaceae did not separate from the other eudicots, even though both groups are known to have unique metabolism, suggesting that more research and curation on members of these groups is needed. This analysis also indicated that despite being represented by a number of PMN species, the unique metabolisms of these groups remain understudied. The separation of Brassicaceae from the other groups may reflect a more comprehensive body of knowledge about the metabolism of *Arabidopsis* due to its status as a model plant and, as a result, a larger number of Brassicaceae-specific pathways being known than for compounds specific to other clades. The same might be true of the grasses, a clade that contains economically important crops such as maize, rice, wheat, and switchgrass. These results suggest that study of representative members of a group could help differentiate the group as a whole and suggest that much of current knowledge is limited to common pathways. More detailed studies of the metabolism of other groups are needed to understand what makes them unique.

### Previous work making use of PMN

PMN has been used in a variety of ways by the plant research community. One common use is to find metabolic information about a specific area of metabolism, such as finding sets of biosynthesis genes for a particular compound or sets of compounds under study, or finding pathways associated with a set of genes highlighted by an experiment. Clark and Verwoerd (2011) used AraCyc to determine different biosynthetic routes for anthocyanin pigments and predict minimal sets of genes which could be mutated to eliminate pigment production. Pant et al. (2015) performed metabolite profiling on phosphorus-deprived *Arabidopsis* wild type plants and phosphorus-signaling mutants. PMN was used to find genes in the biosynthetic pathways of metabolites which showed altered concentration in the mutants and P-deprived plants. Saptari and Susila (2018) examined the expression of hormone biosynthesis genes during somatic embryogenesis in *Arabidopsis* and rice. The authors used PMN to identify hormone biosynthetic genes and performed expression analysis on the identified gene set. Kooke et al. (2019) used AraCyc (alongside other databases) to identify genes involved in glucosinolate and flavonoid metabolism, and then examined the relationship between methylation of these genes and metabolic trait values. Uhrig et al. (2020) examined diurnal changes in protein phosphorylation and acetylation, and used PMN’s pathway enrichment feature to identify AraCyc pathways enriched for proteins associated with these protein modification events.

A second common use of PMN is to study broader patterns in plant metabolism. Hanada et al. (2011) explored two rival hypotheses which attempt to explain the large number of *Arabidopsis* metabolic genes for which single mutants show weak or no phenotypes, and used data from PMN to determine the connectivity of different metabolites in the network. Chae et al. (2014) compared primary and specialized metabolism in plants and green algae and found that specialized metabolism genes have different evolutionary patterns from primary metabolism genes. Moore et al. (2019) used AraCyc in assembling lists of enzyme-coding genes involved in primary and specialized metabolism, and then explored associations between various qualities and metrics of the genes and their involvement in primary or specialized metabolism. The PlantClusterFinder (Schläpfer et al. 2017) software was also used in that analysis. Song et al. (2020) set out to test the hypothesis that stoichiometric balance imposes selection on gene copy number. AraCyc pathways were used as a source of functionally-related gene groups to test for reciprocal retention.

A third use of PMN is in genome annotation. Gupta et al. (2015) used RNA-seq data from blueberry (*Vaccinium corymbosum*) to annotate a draft genome sequence for the plant. Gene models were BLASTed against metabolic genes from AraCyc and other species-specific pathway genome databases, and the results were used to improve the annotations. The annotations were then used to examine blueberry metabolism. Similarly, Najafabadi et al. (2017) took transcriptomes of *Ferula gummosa* Boiss., a relative of carrot that is the source of the aromatic resin galbanum, and used BLASTx against enzyme-coding genes from PMN as a source for annotation of enzyme-coding genes in *Ferula*.

## Conclusions

PMN provides an important resource for organizing and making accessible plant metabolism information. The study of plant metabolism enables improvement of the productivity, nutrition, and resilience of crop plants, and furthers understanding of how wild plants function in their ecosystems. PMN data and tools have been used by researchers to answer a broad range of biological questions from development to physiology to evolution. The latest release of PMN, PMN 15, has the breadth and depth of metabolic information that should enable even a wider spectrum of questions to be pursued in plant biology.

## Methods

### The PMN pipeline

New plant databases introduced in each version of PMN are Tier 3 BioCyc databases (Karp et al. 2019), which indicate that the information is based mostly on automated prediction using their genome. Any experimentally-supported enzymes and pathways in Metacyc or Plantcyc that are annotated as belonging to the organism are also imported into the database along with their citations and codes for the type of evidence the cited papers present. The plant’s remaining complement of enzymes is predicted, and its metabolites and pathways are in turn predicted based on the enzymes.

Bringing a new species or subspecies into PMN begins with the sequenced and annotated genome with predicted protein sequences. To be considered for inclusion, a genome must pass a quality metric in the form of BUSCO (Benchmarking Single-Copy Orthologs) (Simão et al. 2015; Waterhouse et al. 2018), which assesses genome completeness using a database of proteins expected to be present in all eukaryotes, with matches assessed using HMMER (http://hmmer.org) (Potter et al. 2018). A score of at least 75% “complete” is required for inclusion in PMN. If a genome passes this metric, it can then be run through the PGDB creation pipeline. First, splice variants are removed, leaving one protein sequence per gene, with the longest variant being retained. The sequences are classified as enzymes or non-enzymes, and enzymatic functions are predicted, using the Ensemble Enzyme Prediction Pipeline (E2P2) software (Chae et al. 2014; Schläpfer et al. 2017). E2P2 uses BLAST and PRIAM to assign enzyme function based on sequence similarity to proteins with previously-known enzymatic functions based on functional annotations taken from several sources including MetaCyc (Caspi et al. 2020), SwissProt (UniProt Consortium 2021), and BRENDA (Chang et al. 2021). The genomes included in PMN 15 were checked using BUSCO v 3.0.2 using the Eukaryota ODB9 dataset. Enzyme prediction for PMN 15 was done using E2P2 v4.0 and RPSD v4.2, which was generated using data from PlantCyc 12.5, MetaCyc 21.5, BRENDA (downloaded April 4, 2018), SwissProt (downloaded April 4, 2018), TAIR (downloaded April 5, 2018), Gene Ontology (Downloaded April 4, 2018), and Expasy (release of March 28, 2018).

Once enzymes are predicted, they must be assembled into pathways by the PathoLogic function of Pathway Tools (Karp et al. 2019). The set of predicted pathways is then further refined using the Semi-Automated Validation Infrastructure (SAVI) software (Schläpfer et al. 2017). SAVI is used to automatically apply broad curation decisions to the pathways predicted for each species. It can be used, for example, to specify particular pathways that are universal among plants and should therefore be included in all species’ databases even if not predicted by PathoLogic. SAVI can also be used to specify that a particular pathway is known to be present only within a specific plant clade. Therefore, if the pathway is predicted in a species outside of that clade, it should be considered a false prediction and removed. PMN 15 was generated using Pathway Tools 24.0 and SAVI 3.1.

The final parts of the pipeline are grouped into three stages: refine-a, refine-b, and refine-c. In refine-a, the database changes recommended by SAVI are applied to the database and pathways added or approved by SAVI have SAVI citations added. In refine-b, pathways and enzymes with experimental evidence of presence in a plant species are added to that PGDB if they were not predicted, and appropriate experimental evidence codes are added. In refine-c, authorship information is added to the PGDB, the cellular overview is generated, and various automated data consistency checks are run.

The convention for PGDB versions was updated in PMN 15. Taking SorghumbicolorCyc 7.0.1 as an example, the first number, 7, is incremented when the PGDB is re-generated *de novo* from a new version of MetaCyc and/or a new genome assembly. The second, 0, is incremented when there are error corrections or other fixes to the content of the database. A third, 1 in the example, may be added when the database is converted to a new version of Pathway Tools without being regenerated, a process that does not alter the database contents.

### Changes in curation policy

Since its initial 1.0 release, some changes in curation policy have been made to PMN and PlantCyc. In 2013, the *Arabidopsis*-specific database, AraCyc, switched from identifying proteins by locus ID to using the gene model ID. This eliminates ambiguity when multiple splice variants exist for a single locus. In PMN 10, the policy for all species was switched from using the first splice variant to the longest. This was done because a longer splice variant is likely to have more domains, making it easier to determine its function.

In PMN 10, the database narrowed its focus strictly to small-molecule metabolism, and pathways involved solely in macromolecule metabolism (such as protein synthesis) were removed. Macromolecules have never been the focus of PMN, and provision of information about them is a role better served by other databases with tools specifically suited to large heteropolymers like proteins and DNA/RNA.

In version 13 of PMN, the PlantCyc database was limited to only include pathways and enzymes with experimental evidence to support them. The original purpose of including all information, experimental and computational, in PlantCyc was to allow cross-species comparison, a function now served by the virtual data integration and display functionality recently introduced in Pathway Tools (Karp et al. 2019). PlantCyc now serves as a repository of all experimentally-supported compounds, reactions, and pathways for plants.

### Manual pathway prediction validation

120 PMN pathways were randomly selected to manually assess pathway prediction accuracy. The 126 organism-specific PGDBs were then re-generated using E2P2 and PathoLogic alone, with PathoLogic set to ignore the expected phylogenetic range of the pathway and call pathway presence / absence based only on the presence of enzymes (no taxonomic pruning), no SAVI, and skipping the step of importing pathways with experimental evidence of a species into that species database if the pathway was not predicted. This resulted in a set of PGDBs based purely on computational prediction that we refer to as “naïve prediction PGDBs”. Biocurators evaluated the accuracy of each of the 120 pathway’s prediction across all 126 organisms in PMN in the naïve prediction PGDBs and, separately, in the released version of PMN. Specifically, we evaluated whether pathway assignments to the PGDBs reflected the taxonomic range of the pathway as expected from the literature. Each pathway’s assignment to the naïve prediction PGDBs and released PGDBs was classified with respect to the expected taxonomic range as either “Expected” (predicted and expected species are mostly the same), “Broader” (pathway is predicted beyond its expected range), “Narrower” (predicted range of the pathway is smaller than the expected range), or it was identified to be a non-plant or non-algal pathway, and therefore classified as a non-PMN pathway.

### Presence-absence matrices

In order to analyze the pathways, reactions, and compounds (PRCs) in each species’ database, presence-absence matrices were generated for each of these three data types. Each is a binary matrix containing the list of PMN organisms as its rows and a list of PRCs of one type as its columns. Each matrix element is equal to 1 if the organism contains the PRC and 0 if it does not (Supplemental Files S1-S3). Reactions were only marked as present in a species if the species had at least one enzyme annotated to the reaction, whether predicted or from experimental evidence. Since PRCs that are present in either only one organism or all organisms are not useful in differentiating plant groups, we excluded these PRCs from further analysis. Separately, a table was generated that maps the species to one of several pre-defined taxonomic groups (Supplemental File S4). The groups were selected manually to best represent the diversity of species in PMN and included monophyletic and paraphyletic groups, as well as a polyphyletic “catch-all” group (“Other angiosperms”). The PRC matrices and the plant group table were used to investigate relationships among the species through the lens of metabolism. The PRC matrices were produced using a custom lisp function (Supplemental File S5).

### Multiple correspondence analysis

The PRC matrices were used to perform multiple correspondence analysis (MCA) (Greenacre et al. 2006). This is a technique similar to principal component analysis (PCA) but is frequently used with categorical (binomial or multinomial) data. It differs from PCA in that a complete disjunctive table (CDT) is first produced from the input matrix. In a CDT, each multinomial variable i (a column in the input matrix) is split into *J*_*i*_ columns where *J*_*i*_ is the number of levels of variable *i*. In this analysis, the variables are the pathways, reactions, or compounds (PRCs), and there are two levels for each, present and absent. Each CDT column *j*_*i*_ therefore corresponds to one level of one variable and is initially set equal to 1 for species for whom that PRC is present and 0 otherwise. Each group of *J*_*i*_ columns therefore contains, in each row, one column equal to 1 and *J*_*i*_*–1* columns equal to 0. In this analysis, therefore, each pathway results in two columns in the CDT, set to 1 0 if the pathway is present and 0 1 if the pathway is absent. MCA then scales the values of each column in the CDT according to the rarity of that level of that variable, so that each CDT column sums to 1. The remainder of the procedure is the same as in PCA. Because of the scaling, a species will be further from the origin in the MCA scatterplot if it possesses uncommon PRCs or lacks common ones. The MCA was performed using the MCA() function of the R package FactoMineR v2.3 (Lê et al. 2008). The MCA scatter plots were colored using the columns of the plant group table (Supplemental File S4) to elucidate relationships between the MCA clusters and plant groups. The scatter plots were generated using ggplot2 v3.3.4.

### Metabolic domain enrichment

To examine the pathways associated with each MCA axis, the percentage of variance explained by the presence or absence of each pathway, found in pwy.mca$var$contrib (where pwy.mca is the R object returned by FactoMineR’s MCA function when run on the pathway matrix), was exported to a tab-delimited text file. To determine which metabolic domains, if any, were overrepresented in the set of pathways describing the variance of MCA dimensions 1 and 3, we ran an enrichment analysis of the set of pathways explaining the 95^th^ percentile of the variance. Pathways were mapped to a metabolic domain using supplementary information from (Schläpfer et al. 2017). Pathways left unmatched were manually assigned to a metabolic domain by expert curators and a new pathway-metabolic domain mapping file version 2.0 was created (Supplemental Table S7). Enrichment background was set as all pathways from PMN’s 126 organism-specific databases, all of which were assigned to metabolic domains. Enrichment was calculated using the phyper() function from the R stats package and p-values were corrected for multiple hypothesis testing at a false discovery rate (FDR) of 5%.

### Omics Dashboard and Enrichment Analysis

The sorghum drought transcriptomics data from (Varoquaux et al. 2019) were downloaded from: https://www.stat.berkeley.edu/~epicon/publications/rnaseq/. We specifically used their log-fold change and differential expression analysis results. For both leaf and root samples, the sets of all expressed genes were filtered to include only those differentially expressed in either cultivar as a result of post-flowering drought (using an FDR of 5%). Corresponding expression data for both gene sets were then filtered to include only the week prior to, and the first two weeks of post-flowering drought (weeks 9-11). The resulting data sets were then directly uploaded into PMN’s Omics Dashboard for visual analysis of metabolic trends. Enrichment analysis of metabolic genes among leaf and root DEGs as a result of post-flowering drought was calculated in R version 3.6.3 with a hypergeometric test using the phyper() function from the stats package. The background used for this enrichment analysis was all *Sorghum bicolor* genes (McCormick et al. 2018) from the *Sorghum bicolor* genome annotation v3.1.1 downloaded from Phytozome v12. Violin plots were generated using the geom_violin() function within the ggplot2 package in R version 3.6.3. Statistical differences between non-metabolic and metabolic DEGs as a function of time were determined by two-way ANOVA followed by Tukey’s Honest Significant Difference (HSD) test (p < 0.05) using the lsmeans() functions within the lsmeans package in R version 3.6.3. Pathway enrichment among the set of metabolic root DEGs was calculated using the “Genes Enriched for Pathways” functionality within the “Enrichments” dropdown of a SmartTable. We performed an enrichment analysis using Fisher’s Exact test and Benjamini-Hochberg correction at an FDR of 5% with the set of all pathway genes from SorghumbicolorCyc (version 7.0.1) as the background.

### Cell type activity analysis

We downloaded and integrated datasets from 5 existing *Arabidopsis* root single-cell RNAseq studies. Briefly, raw fastq files for 21 datasets derived from studies by (Zhang et al. 2019), (Jean-Baptiste et al. 2019), (Denyer et al. 2019), (Ryu et al. 2019), and (Shulse et al. 2019) were downloaded, trimmed, and mapped using the STARsolo tool v.2.7.1a. Whitelists for each dataset were obtained either from the 10X Cellranger software tool v. 2.0 for the 10X-Chromium samples, or after following the Drop-seq computational pipeline (https://github.com/broadinstitute/Drop-seq/releases/tag/v2.3.0), extracting error-corrected barcodes from the final output for the Drop-seq samples. Valid cells within the digital gene expression matrices computed by STARSolo were then determined as those having total unique molecular identifier (UMI) counts greater than 10% of the 1^st^ percentile cell, after filtering for cells with very high (20,000) UMIs. Cells containing greater than 10% mammalian reads, greater than 10% organellar (chloroplast or mitochondrial) reads, or cells having transcripts from fewer than 200 genes were filtered out. Filtered digital gene expression matrices were then pre-processed using the Seurat (v3.1.0) package after removing protoplast-inducible genes (Birnbaum et al. 2003), using the SCTransform method (with 5000 variable features). All Seurat objects were then integrated together using the approach from (Stuart et al. 2019), applying the SelectIntegrationFeatures, PrepSCTIntegration, FindIntegrationAnchors, and IntegrateData functions from the Seurat R package, using 5000 variable features, 20 principal components, and otherwise default parameters. Cell clusters were computed using the Seurat functions, FindNeighbors and Find Clusters, 20 principal components and a resolution parameter of 0.8. Index of Cell Identity (ICI) scores were computed using a combination of existing ATH1 microarray and RNAseq single cell datasets (Supplemental Table S6). Briefly, arrays were normalized using the gcrma R package, and RNA-seq data were trimmed using the bbduk tool, and mapped using bbmap (sourceforge.net/projects/bbmap/). Transcript counts were quantified using the featureCounts tool (Liao et al. 2014). Raw RNAseq counts were then normalized using the edgeR package (v 3.26.0), with the “upperquartile” method. Normalized reads were then further normalized with the gcrma-normalized microarray data using the Feature-Specific Quantile Normalizations (FSQN) method (Franks et al. 2018) to obtain a dataset consisting of both RNA-seq and microarray-based cell-type specific transcriptome measurements. This dataset was then used to build an ICI (Birnbaum and Kussell 2011) specification matrix using the methods described by (Efroni et al. 2015). This specification table was then used to compute ICI scores for each cell in the integrated single-cell dataset, along with p-values derived from random permutation.

To map the single-cell data to metabolic domains, pathways, and enzymes, we used AraCyc v.17.0, which includes 8556 metabolic genes and 650 pathways. We used the pathway-metabolic domain mapping file version 2.0 (Supplemental Table S7) to map the pathways to 13 metabolic domains. To avoid biases introduced by small sample size to the cell type specificity analysis, we only included pathways containing at least 10 genes whose transcripts were detected in the single cell data described above. Based on these criteria, 198 out of 650 pathways were included in this analysis. To compute cell type specificity at the transcript level, we first calculated the expression level for a pathway or domain per cell type by taking the average of expression values for all the genes annotated to this pathway or domain within this cell type. The cell type specificity was defined as the cell type(s) for which the expression level of a pathway or domain was at least 1.5-fold higher than their background expression, which was calculated by taking the average of expression values for all the genes annotated to this pathway or domain in all cells. Since the expression levels of a pathway or domain per cell type could be influenced by gene expression outliers, we only included the cell types in which more than 50% of genes associated with the pathway or domain showed higher expression than their background expression based on a Wilcoxon test followed by a multiple hypothesis test adjustment using FDR with a threshold of 0.01. The background expression level of a gene was calculated by taking the average of its expression values in all the cells included in this study. Heatmaps were generated using the R package ggplot2 v.3.1. To compute cell type specificity at the pathway level, we first selected the set of pathways containing at least 10 genes whose transcripts were captured by the single cell transcriptomic data to avoid biases that could be introduced by small sample size. Based on these criteria, 30% (198 out of 650) *Arabidopsis* pathways were included in this analysis.

In a metabolic network, isozymes are defined as enzymes encoded by different genes catalyzing the same reaction, which are usually the result of gene duplication events. To investigate whether isozymes tend to be expressed in different cells compared to enzymes catalyzing different reactions within the same pathway, we analyzed gene expression pattern similarity between a pair of enzymes across *Arabidopsis* root cells by computing Spearman’s correlation. To prevent having correlations between self, we removed enzymes that are mapped to more than one reaction in a pathway as well as pathways that contain only one reaction. Spearman’s correlation coefficients were computed using the function cor() in R. Significant correlation coefficients were determined using an R package scran v.1.18.5 (Lun et al. 2016). Distribution of Spearman’s rho was compared using a one-way ANOVA followed by post-hoc adjustment with Tukey’s test in R. The box plot was generated using the R package ggplot2 v.3.1.

## Author Contributions

S.Y.R. conceived the project. C.H., A.X., and B.X. developed the pipelines and generated PMN databases. D.G., S.R., and W.D. evaluated the quality of the databases. C.H., S.R., and W.D. compared the databases using MCA analysis. D.G. performed Omics Dashboard analysis using sorghum drought transcriptome data. K.Z. analyzed PMN’s AraCyc data using *Arabidopsis* root single cell-type transcriptome data. B.C. curated the *Arabidopsis* root single cell-type transcriptome data. B.X. developed the Co-Expression Viewer. S.P. and P.K. developed the Pathway Tools software, including the subcellular compartmentalization viewer. A.X. drew plant artwork for Figure 1. C.H. wrote the manuscript with contributions from D.G., K.Z., W.D., B.C., and S.Y.R. All authors edited the manuscript. S.Y.R. supervised the project and manuscript preparation. S.Y.R. agrees to serve as the author responsible for contact and ensures communication.

## Acknowledgements

We thank members of the Rhee lab for helpful discussions. This work was supported by grants from the National Science Foundation (IOS-1546838, IOS-1026003) and the U.S. Department of Energy, Office of Science, Office of Biological and Environmental Research, Genomic Science Program grant nos. DE-SC0018277, DE-SC0008769, DE-SC0020366, and DE-SC0021286. We thank Justin Krupp and Jason Thomas for editing the manuscript. We also thank Brenda Yu for her work in constructing the 45k cell dataset. The gymnosperm illustration in Figure 1C was created with BioRender.com.

## Abbreviations used

ANOVA: analysis of variance
BUSCO: Benchmarking Single-Copy Orthologs
CDT: complete disjunctive table
ChEBI: Chemical Entities of Biological Interest
E2P2: Ensemble Enzyme Prediction Pipeline
GO: Gene Ontology
JI: Jaccard Index
KEGG: Kyoto Encyclopedia of Genes and Genomes
MCA: multiple correspondence analysis
NPP: Non-PMN pathway
PCA: principal component analysis
PGDB: pathway genome database
PMN: Plant Metabolic Network
PRC: pathway, reaction, or compound
SAVI: Semi-automated Validation Infrastructure

